# In-Depth Mapping of DNA-PKcs Signaling Uncovers Conserved Features of Its Kinase Specificity

**DOI:** 10.1101/2024.01.17.576037

**Authors:** Shannon Marshall, Marcos V.A.S. Navarro, Carolline F.R. Ascenҫão, Marcus B. Smolka

**Affiliations:** Department of Molecular Biology and Genetics, Weill Institute for Cell and Molecular Biology, Cornell University, Ithaca, NY 14853, USA; IFSC Institute of Physics of São Carlos, University of São Paulo, São Carlos - SP, 13566-590, Brazil

## Abstract

DNA-PKcs is a DNA damage sensor kinase with established roles in DNA double-strand break repair via non-homologous end joining. Recent studies have revealed additional roles of DNA-PKcs in the regulation of transcription, translation and DNA replication. However, the substrates through which DNA-PKcs regulates these processes remain largely undefined. Here we utilized quantitative phosphoproteomics to generate a high coverage map of DNA-PKcs signaling in response to ionizing radiation and mapped its interplay with the ATM kinase. Beyond the detection of the canonical S/T-Q phosphorylation motif, we uncovered a non-canonical mode of DNA-PKcs signaling targeting S/T-ψ-D/E motifs. Cross-species analysis in mouse pre-B and human HCT116 cell lines revealed splicing factors and transcriptional regulators phosphorylated at this novel motif, several of which contain SAP domains. These findings expand the list of DNA-PKcs and ATM substrates and establish a novel preferential phosphorylation motif for DNA-PKcs that connects it to proteins involved in nucleotide processes and interactions.

## INTRODUCTION

DNA-dependent Protein Kinase (DNA-PK) is a Phosphatidylinositol 3-Kinase-related Kinase (PIKK) with key roles in the repair of DNA double strand breaks (DSBs) through non-homologous end joining (NHEJ) (Davis *et al*, 2014). DNA-PK is a holoenzyme composed of a large catalytic subunit, DNA-PKcs, and the Ku70/Ku80 heterodimer (Dvir *et al*, 1992; Gottlieb & Jackson, 1993). DNA-PK is rapidly recruited to broken DNA ends by Ku, which induces conformational changes that activate the catalytic activity of DNA-PKcs (Chen *et al*, 2021; Buehl *et al*, 2023). While active DNA-PKcs is reported to phosphorylate hundreds of proteins, including several components of the NHEJ complex such as Ku, XRCC4, and XLF (Kurosawa, 2021), the best characterized substrate currently demonstrated to be required for NHEJ is DNA-PKcs itself (Ding *et al*, 2003; Meek *et al*, 2008). DNA-PKcs autophosphorylation alters the conformational state of DNA-PKcs engaged at DNA ends, granting access of broken DNA ends to processing enzymes like the nuclease Artemis and the Pol X family polymerases, in addition to stimulating the release of DNA-PKcs from DNA ends (Liu *et al*, 2022). Paradoxically, autophosphorylation seems to also limit the extent to which DNA ends can be processed (Cui *et al*, 2005). Loss of DNA-PKcs results in severe DNA repair defects, which manifest at the organismal level as severe combined immunodeficiency (SCID) since V(D)J and class switch recombination require DSB end processing and joining via NHEJ (Blunt *et al*, 1995; Kirchgessner *et al*, 1995; Liu *et al*, 2022).

Apart from its canonical role in DNA repair, DNA-PKcs has also been implicated in a range of other nuclear processes. For example, DNA-PKcs promotes faithful replication of telomeres and telomere capping through phosphorylation of hnRNPA1 (Le *et al*, 2013; Sui *et al*, 2015; Flynn *et al*, 2011). Additionally, DNA-PKcs has been shown to phosphorylate and regulate various transcription factors. In fact, one of its first identified substrates was the transcription factor SP1 during the formation of promoter bound transcriptional complexes (Jackson *et al*, 1990). Inhibition or depletion of DNA-PKcs was reported to result in reduction in RNA Pol II-mediated transcription in a manner that is dependent on the DNA-PKcs substrate TRIM28 (Bunch *et al*, 2015). The catalytic activity of DNA-PKcs is also relevant for ribosome biogenesis. DNA-PKcs makes Ku-dependent contacts with the snoRNA in the U3 component of the ribosomal small subunit processome, where it is activated and autophosphorylated (Shao *et al*, 2020). Mutations in DNA-PKcs that impair its catalytic activity or block autophosphorylation halt rRNA processing, resulting in ribosome deficiency and translation defects in haematopoietic cells. More recently, DNA-PKcs has been shown to be involved in the control of DNA replication in response to replication stress by promoting replication fork reversal and the slow-down of fork progression (Dibitetto *et al*, 2022).

The precise targets through which DNA-PKcs mediates its functions in NHEJ-mediated DNA repair, RNA processing and DNA replication remain largely unknown. Identification of functional substrates is complicated by the lack of a complete map of DNA-PKcs signaling and by the partial redundancy of its roles and substrates with the ATM kinase. Here we performed phosphoproteomic analysis of DNA-PKcs signaling and mapped its division of labor with the ATM kinase. To gain deep coverage of the DNA-PKcs and ATM signaling network, we used Lig4-deficient mouse Pre-B cells that are unable to efficiently repair DSBs, and therefore hyper-accumulate DNA-PKcs and ATM signaling induced by ionizing radiation (IR). Our experimental setup allowed us to define the unique contributions of ATM and DNA-PKcs kinases and uncover a novel S/T-ψ-D/E motif preferentially targeted by DNA-PKcs in substrates involved in RNA processing and transcription. We found that the U2 snRNP splicing factor SF3A3 is directly phosphorylated by DNA-PKcs at an S/T-ψ-D/E site, which alters the affinity of SF3A3’s SAP domain towards double-stranded DNA. These findings expand the list of DNA-PKcs and ATM substrates and establish a novel preferential phosphorylation motif for DNA-PKcs that connects it to proteins involved in RNA biology.

## RESULTS

### In-depth mapping of the signaling response induced by ionizing radiation (IR) in mouse pre-B cells

We conducted a phosphoproteomic analysis of IR-induced phosphorylation using mouse pre-B cells transformed with Abelson murine leukemia virus (A-MuLV) and deficient for DNA Ligase IV (LIG4) (Bredemeyer *et al*, 2006). This system provides several advantages for mapping DNA-PKcs signaling with high coverage (Figure 1A). First, these pre-B cells can be robustly arrested in G1 using treatment with the ABL kinase inhibitor imatinib, helping to minimize confounding effects from ATR activation in S-G2 (Wilson *et al*, 2010; Muljo & Schlissel, 2003). Second, Abelson Pre-B cell nuclei account for approximately 70% of their cellular volume (Ulloa *et al*, 2021), which enhances the coverage of nuclear proteins in our phosphoproteomics analysis. Third, these cells are deficient in DNA ligase IV, a critical enzyme for the repair of DSBs through NHEJ, which is the predominant repair mechanism during the G1 phase. This will result in an accumulation of unrepaired breaks and persistence of DSB signaling (Rothkamm *et al*, 2003).

**Figure 1.**
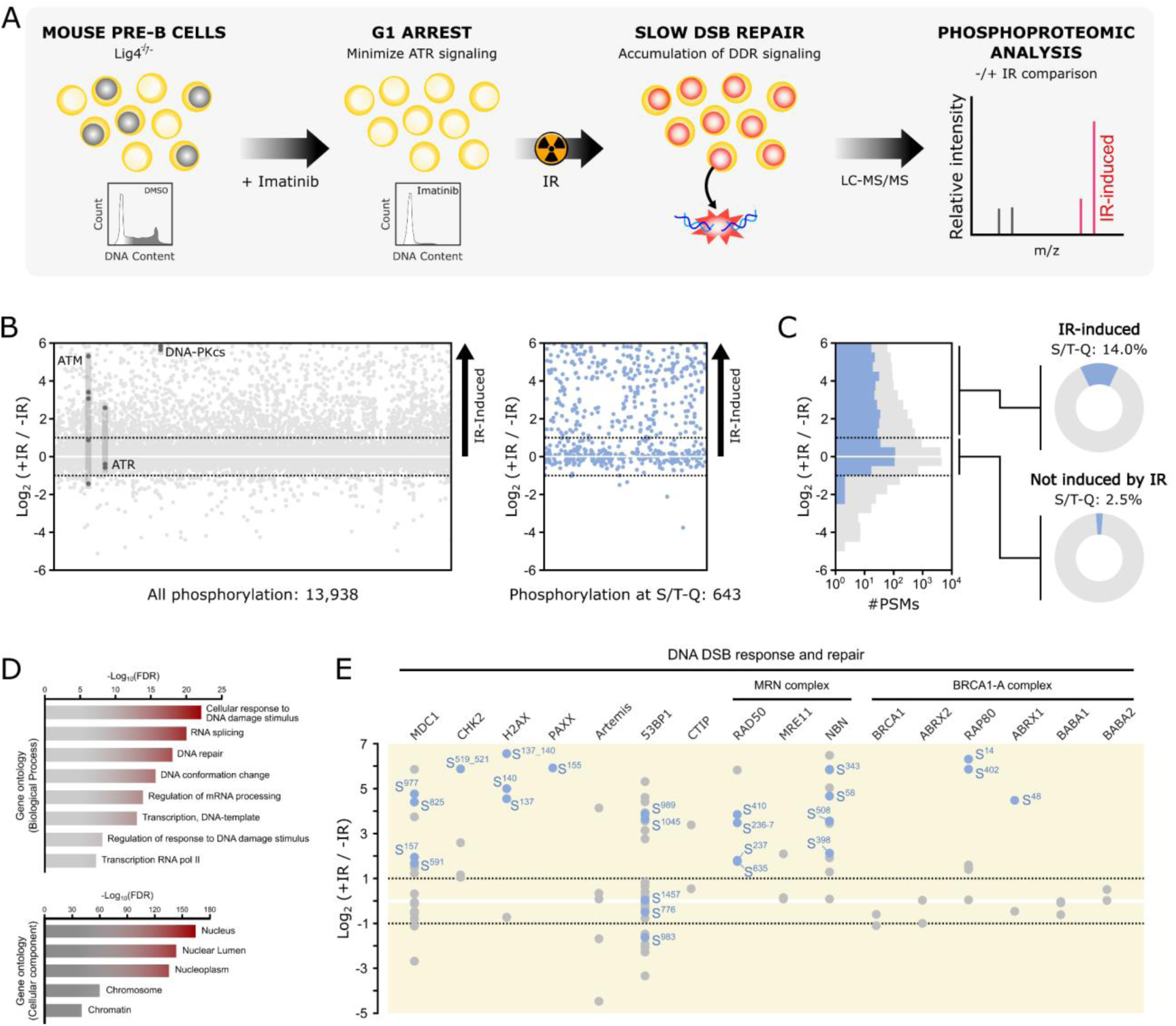
In-depth phosphoproteomic analysis of the cellular response to IR in mouse pre-B cells. A) Workflow for maximizing the detection of IR-induced phosphorylation in mouse pre-B cells. Pre-B cells have large nuclei, and high nuclear to cytosolic protein ratio, which facilitates detection of low abundance nuclear signaling events. Deletion of LIG4 prevents the rapid repair of IR-induced breaks, leading to accumulation of IR-induced signaling events. Pre-B cells from Lig4^-/-^ mice were first arrested in G1 with imatinib. Following arrest, cells were irradiated with 20 gy IR and harvested for quantitative phosphoproteome analysis. Quantification of phosphoproteomic changes was accomplished using SILAC. B) The plot on the left compares the phosphoproteome of pre-B cells treated with IR to the phosphoproteome of control (untreated) cells. The plot on the right displays S/T-Q sites only. Dashed lines indicate a 2-fold change in abundance. Sites identified in ATR, ATM and DNA-PKcs are highlighted. Each point represents the average of at least two independent experiments (see Supp. Figure 1). C) Cumulative plot of S/T-Q sites (blue) versus all sites (gray). Pie charts on the left highlight the proportion of S/T-Q sites not induced (-1 < Log_2_ (+IR / -R) < 1) and induced (Log_2_ (+IR / -R) > 1) by IR. D) Curated gene ontology analysis showing enriched biological processes and cellular components among IR induced phosphorylation sites. E) Selected phosphorylation sites identified in proteins involved in DNA double strand break response and repair. Dashed lines indicate a 2-fold change in abundance. S/T-Q sites are light-blue.

G1-arrested cells were treated with 20 gy of IR and then harvested 90 minutes later for analysis using SILAC-based mass spectrometry. Over 16,000 phosphorylation sites were identified, mapping to over 3,900 proteins (Supp. Table 1). To ensure confidence in our dataset, we utilized a bowtie filtering approach based on reciprocal labeling (Faca *et al*, 2020), resulting in a total of 13,938 high-quality phosphopeptide identifications and quantifications (Figure 1B; Supp. Figure 1). Among these phosphopeptides, 2,258 exhibited at least a 2-fold increase in abundance after IR treatment and were categorized as IR-induced. To our knowledge, this represents the most comprehensive set of IR-induced phosphorylation events reported in mammalian cells (Sampadi *et al*, 2022; Bensimon *et al*, 2010; Bennetzen *et al*, 2010; Winter *et al*, 2017; Wiechmann *et al*, 2020; Yang *et al*, 2010). As expected, the preferential phosphorylation motif for ATM and DNA-PKcs phosphorylation, phosphoS/T-Q, was enriched in the set of IR-induced sites (Figure 1B). Moreover, we detected phosphorylation sites in ATM and DNA-PKcs that were highly induced by IR (Figure 1C), consistent with the expectation that these two PIKKs are highly engaged in the IR-induced response in G1-arrested pre-B cells. Gene ontology analysis indicated that the set of IR-induced phosphorylation events was enriched for proteins involved in DNA damage response and repair, mRNA-related processes, with nuclear and chromatin proteins being highly represented (Figure 1D). Proteins with established roles in the DNA DSB response displayed several IR-induced phosphorylation events, with a predominance of S/T-Q phosphorylation sites, but with other non-S/T-Q sites also detected (Figure 1E). Taken together, these results highlight the effectiveness of our approach in providing a comprehensive dataset of IR-induced phosphorylation. Since the pre-B cells used are efficiently arrested in G1, this dataset is likely to be over-representing targets of DNA-PKcs and ATM and limiting the contributions of ATR to the generated signaling responses.

### Mapping ATM and DNA-PKcs-dependent phosphorylation following IR

The significant redundancy among PIKKs in the response to DNA damage presents a major challenge in pinpointing specific kinase targets within DDR signaling pathways (Jette & Lees-Miller, 2015). Leveraging our strategy to effectively minimize ATR signaling, we examined the individual and combined contributions of ATM and DNA-PKcs to the overall IR-induced signaling through selective kinase inactivation using the DNA-PKcs inhibitor (DNA-PKi) NU7441 and the ATM inhibitor (ATMi) KU55933 (Leahy *et al*, 2004; Zhao *et al*, 2006; Ivanov *et al*, 2009; Hickson *et al*, 2004). Inhibition of DNA-PKcs impaired 145 sites, which represents ∼7% of all the IR-induced phosphorylation sites detected in the analysis (Figure 2A). Individual inhibition of ATM impaired 185 sites, which represents ∼19.5% of all the IR-induced phosphorylation sites detected in this specific analysis (Figure 2B). Simultaneous inhibition of DNA-PKcs and ATM caused a reduction in 929 sites, which represents ∼41.1% of the IR-induced phosphorylation sites detected in the experiment in Figure 2C. These results underscore the redundancy of these kinases, where one kinase can largely compensate for the loss of activity of the other kinase. The drastic reduction of IR-induced phosphorylation in the absence of ATM and DNA-PKcs activity is consistent with the prevailing view that these kinases are the major responders to IR-induced DNA damage in G1. About 90% of peptides phosphorylated at the preferred S/T-Q consensus motif were reduced upon dual inhibition of ATM and DNA-PKcs (Supp. Figure 2).

**Figure 2.**
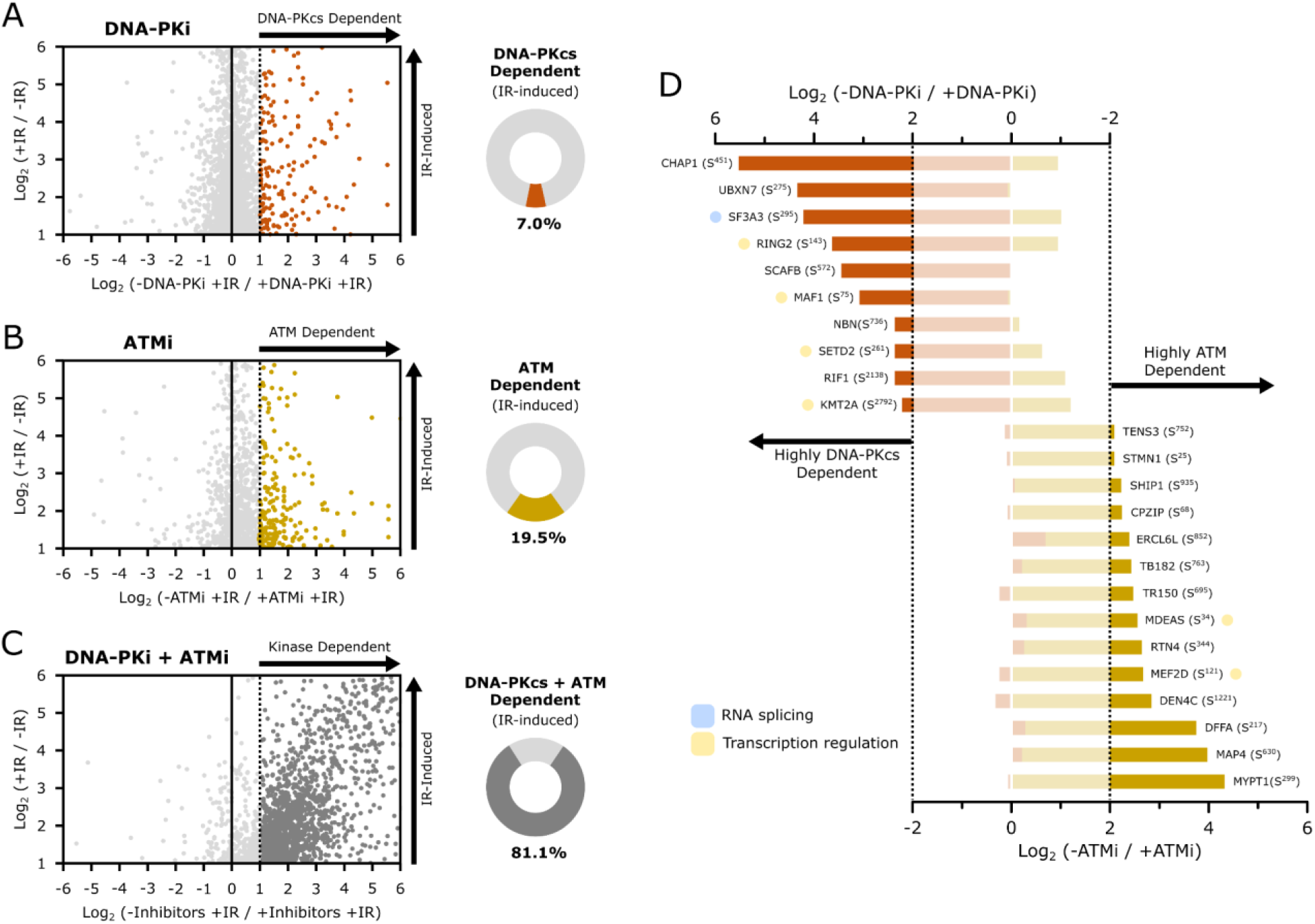
Mapping ATM- and DNA-PKcs-dependent signaling following IR treatment. A-C) Scatter plots comparing the phosphoproteomes of IR-treated Mouse pre-B cells in the absence (y-axis) and presence (x-axis) of standalone DNA-PKi (A) or ATMi (B), and combined kinase inhibitors (C). The pie charts to the right of the scatter plots illustrate the proportions of IR-induced phosphorylation sites dependent on DNA-PKcs and ATM catalytic activities. D) Phosphorylation sites highly dependent on either DNA-PKcs (orange) or ATM (yellow). Dashed lines indicate 4-fold abundance change. Proteins involved in RNA splicing and transcription regulation are highlighted in light blue and light yellow, respectively.

Despite the extensive overlap between ATM and DNA-PKcs dependent substrates in the IR-induced response, our data revealed phosphorylation sites predominantly dependent on each of these kinases, implying a degree of substrate specificity unique for ATM and DNA-PKcs (Figure 2D). For instance, while the phosphorylation status of several spliceosome core components and proteins associated with splicing is influenced by both ATM and DNA-PKcs (Supp. Figure 3), phosphorylation of SF3A3 S^295^ – a core component of the U2 snRNP (van der Feltz & Hoskins, 2019) – is highly dependent on DNA-PKcs but not ATM (Figure 2D). Furthermore, our data reveals differential targeting of factors involved in chromatin remodeling and modification, with the phosphorylation of factors like RING2 and SETD2 being highly dependent only on DNA-PKcs, and MDEAS and MEF2D proteins undergoing phosphorylation only in an ATM-dependent manner (Figure 2D). In addition to their specific contributions to IR-induced signaling, inhibition of both kinases reduced the phosphorylation status of a range of proteins involved in RNA and chromatin biology (Supp. Figure 3). Overall, our data reveal the division of labor for ATM and DNA-PKcs kinases, allowing the delineation of IR-induced signaling into distinct groups based on their level of dependency towards each or both kinases.

### Identification and validation of a novel preferential motif for DNA-PKcs phosphorylation

We analyzed the distribution of phosphorylation motifs in our datasets to identify signaling attributes unique to ATM or DNA-PKcs. We focused on residues at the +1 and +2 positions following the phosphorylation site, and calculated the proportion of amino acids at each position for the groups of sites not induced by IR (Figure 3A) and sites induced by IR displaying dependency for ATM only, DNA-PKcs only or both ATM and DNA-PKcs (Figure 3B). Among the group of phosphorylation events not induced by IR, sites with a proline at the position +1 dominated, consistent with other phosphoproteomic datasets (Lundby *et al*, 2012; Sims *et al*, 2022; Malik *et al*, 2008; Huttlin *et al*, 2010) and expected given the well established action of cyclin-dependent kinases and mitogen-activated protein kinases in cellular homeostasis (Gonzalez *et al*, 1991; Songyang *et al*, 1994). Conversely, the phosphorylation sites with a glutamine at the +1 position (and any amino acid at the +2 position) accounted for only 2.5% of the sites not-induced by IR (Figure 3A). Analyses of IR-induced sites dependent on ATM and DNA-PK, either individually or jointly, unveiled a substantial 6.5- to 10-fold enrichment for the S/T-Q motif, with a slight bias toward negative residues in the +2 position (Figure 3B-C). Strikingly, the set of phosphorylation sites dependent on DNA-PKcs (and not dependent on ATM) displayed a strong enrichment of a phosphorylation motif containing a bulky hydrophobic residue (ψ - F, I, L, or V) at +1 position, and an acidic residue (D or E) at the +2 position (S/T-ψ- D/E motif) (Figure 3B-C). While phosphorylation events at this motif were marginally enriched in the set of ATM-dependent sites, they account for 17.5% of the DNA-PKcs-dependent sites, representing a 4.5-fold increase compared to the frequency observed in the set of phosphorylation events not induced by IR (Figure 3B-C).

**Figure 3.**
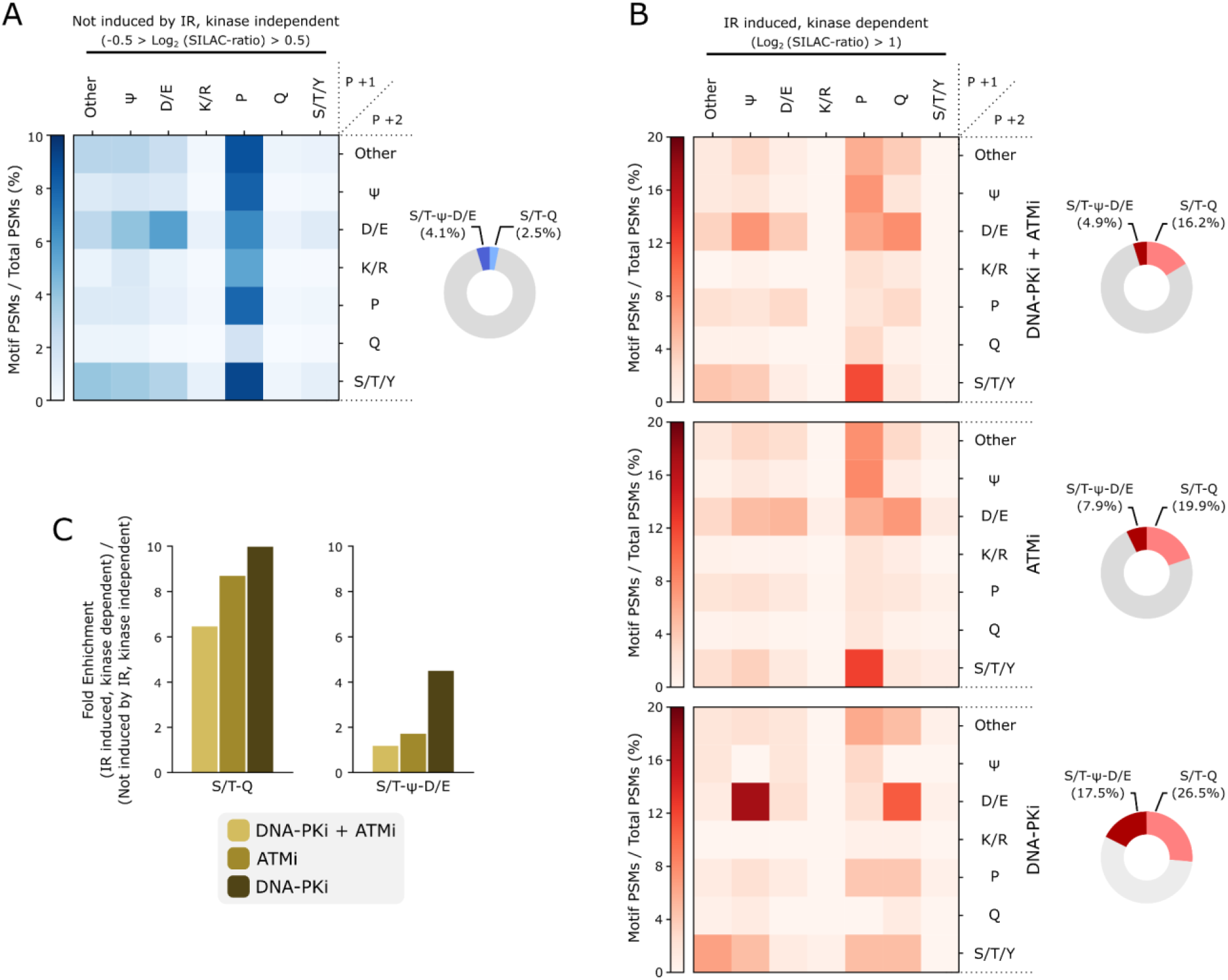
Motif analysis of IR-induced and kinase-dependent phosphorylation sites. A) Heatmap displaying phosphorylation motifs not induced by ionizing radiation (IR) and independent of ATM and DNA-PKcs kinase activity (-0.5 < Log_2_ (SILAC-ratio) < 0.5). B) Heatmaps showing IR-induced phosphorylation motifs dependent (Log_2_ (SILAC-ratio) >1) on the activity of DNA-PKcs (bottom), ATM (center), and both kinases (top). C) Bar charts illustrating the enrichment of S/T-Q and S/T-ψ-D/E motifs that are IR-induced and kinase-dependent relative to non-changing phosphorylations. Motifs are categorized based on the residues at the +1 and +2 positions following the phosphorylated residue. Pie charts to the left of the heatmaps depict the relative proportions of S/T-Q and S/T-ψ-D/E motifs.

To extend these findings and determine whether the S/T-ψ-D/E motif is directly targeted by DNA-PKcs, we used an *in vitro* assay using the purified kinase domain of DNA-PKcs, commercially available, and a chimeric polypeptide substrate we designed, which contained multiple sequences of phosphopeptides we detected to contain a DNA-PKcs-dependent phosphorylation at an S/T-ψ-D/E motif (Figure 4A). From our dataset of DNA-PKcs-dependent phosphorylation, we selected ten phosphopeptides harboring the S/T-ψ-D/E motif and used a sequence of 11 residues surrounding the phosphorylation site (Figure 4B). We cloned the ten sequences in tandem into an *E. coli* expression vector. Following expression and purification, this chimeric substrate was incubated with active recombinant DNA-PKcs, and the resulting phosphopeptides were detected using LC-MS/MS. Our results show that DNA-PKcs can directly phosphorylate all of the ten selected peptides at the S/T-ψ-D/E motif (Figure 4B-C). Notably, mutating the bulky hydrophobic residue in the +1 position or the negative residue in the +2 position to an alanine severely reduced phosphorylation events (Figure 4D), consistent with the S/T-ψ-D/E motif being a preferential and specific motif for DNA-PKcs phosphorylation.

**Figure 4.**
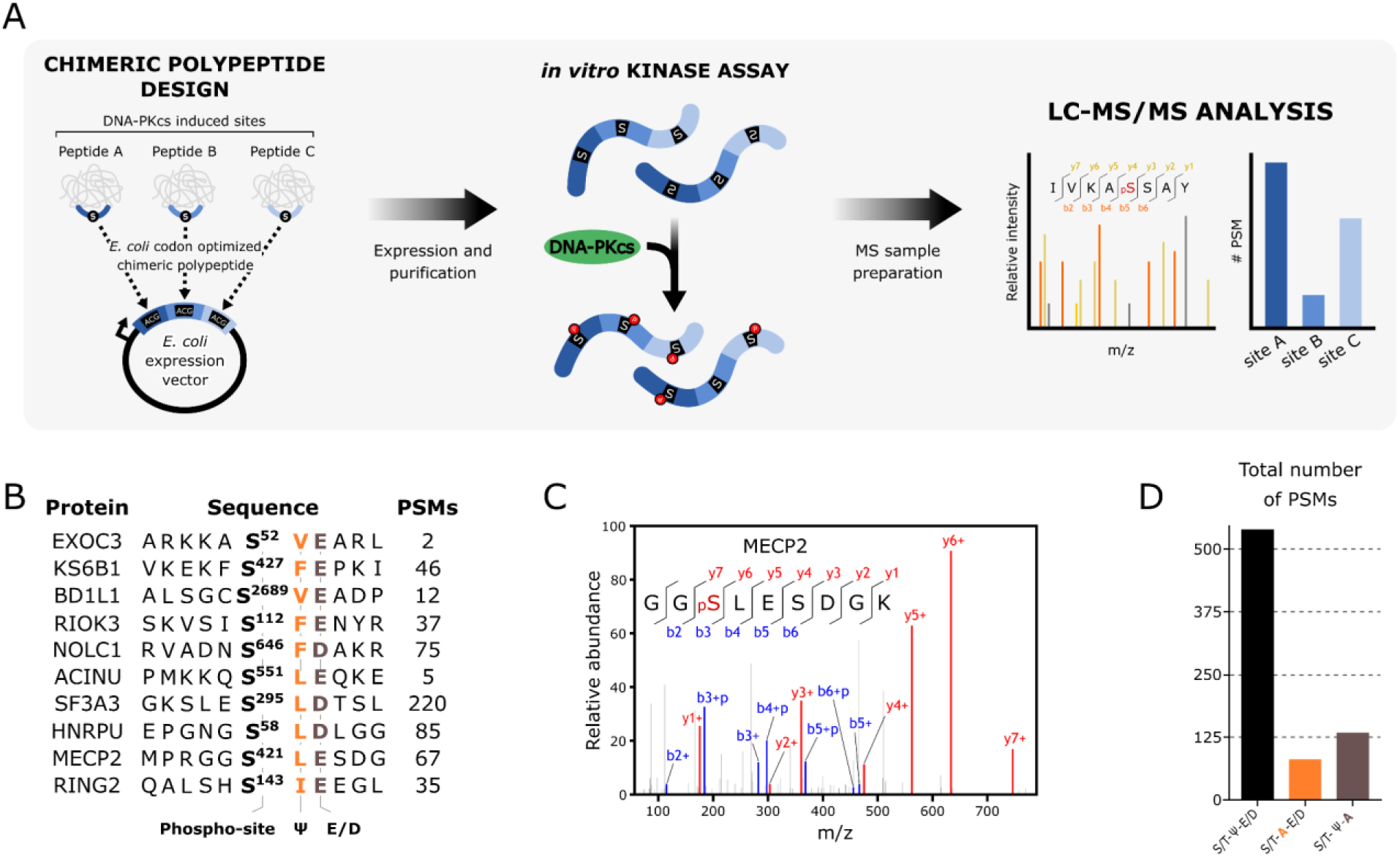
DNA-PK *in vitro* kinase assay validates phospho-targets as direct substrates. A) Schematic workflow of the *in vitro* kinase assay. B) List of proteins and respective peptide sequences included in the chimeric peptide construct. The PSMs column shows the frequency of identification of the phosphorylated peptide post-incubation with DNA-PK. C) Representative MS2 spectrum confirming the identification of the phosphorylated peptide derived from MECP2, encoded within the synthetic chimeric peptide. D) Bar chart depicting the number of PSMs for each phosphorylation site identified in the chimeric peptide construct during the MS experiments, where S/T-A-E/D and S/T-ψ-A indicate constructs containing mutations at the +1 or +2 positions to alanine.

### Specificity towards S/T-ψ-D/E motifs is a conserved feature of DNA-PKcs kinase activity

To evaluate the conservation of DNA-PKcs signaling and its specificity for the S/T-ψ-D/E motif beyond mouse pre-B cells, we carried out a phosphoproteomics analysis of IR-treated asynchronous human HCT116 cells, comparing cells mock-treated or treated with DNA-PKcs inhibitor. Similar data processing as our prior pre-B cell experiment revealed 11,438 phosphorylation sites, with 269 showing at least a 2-fold reduction upon DNA-PKcs inhibition (Supp. Figure 4 and Supp. Table 2).

Using the DIOPT Ortholog Finder (Madeira *et al*, 2022), we identified homologous proteins between human and mouse datasets, and aligned respective phosphorylation sites present in our mouse and human phosphoproteomic datasets. This analysis unveiled over 1,200 conserved phosphorylation sites in mouse pre-B and human HCT116 cells (Supp. Table 3). Figure 5A displays the effect of DNA-PKi on these phosphorylation sites, showing that 31 sites (28 unique proteins) were consistently DNA-PKcs-dependent in both species, including three phosphorylations at the canonical S/T-Q and two at the S/T-ψ-D/E motif. Notably, proteins with a conserved phosphorylation at the S/T-ψ-D/E motif included HNRPU and SF3A3, which were directly phosphorylated by DNA-PKcs *in vitro* (Figure 4C). These results establish S/T-ψ-D/E as a conserved preferential phosphorylation motif for DNA-PKcs.

**Figure 5.**
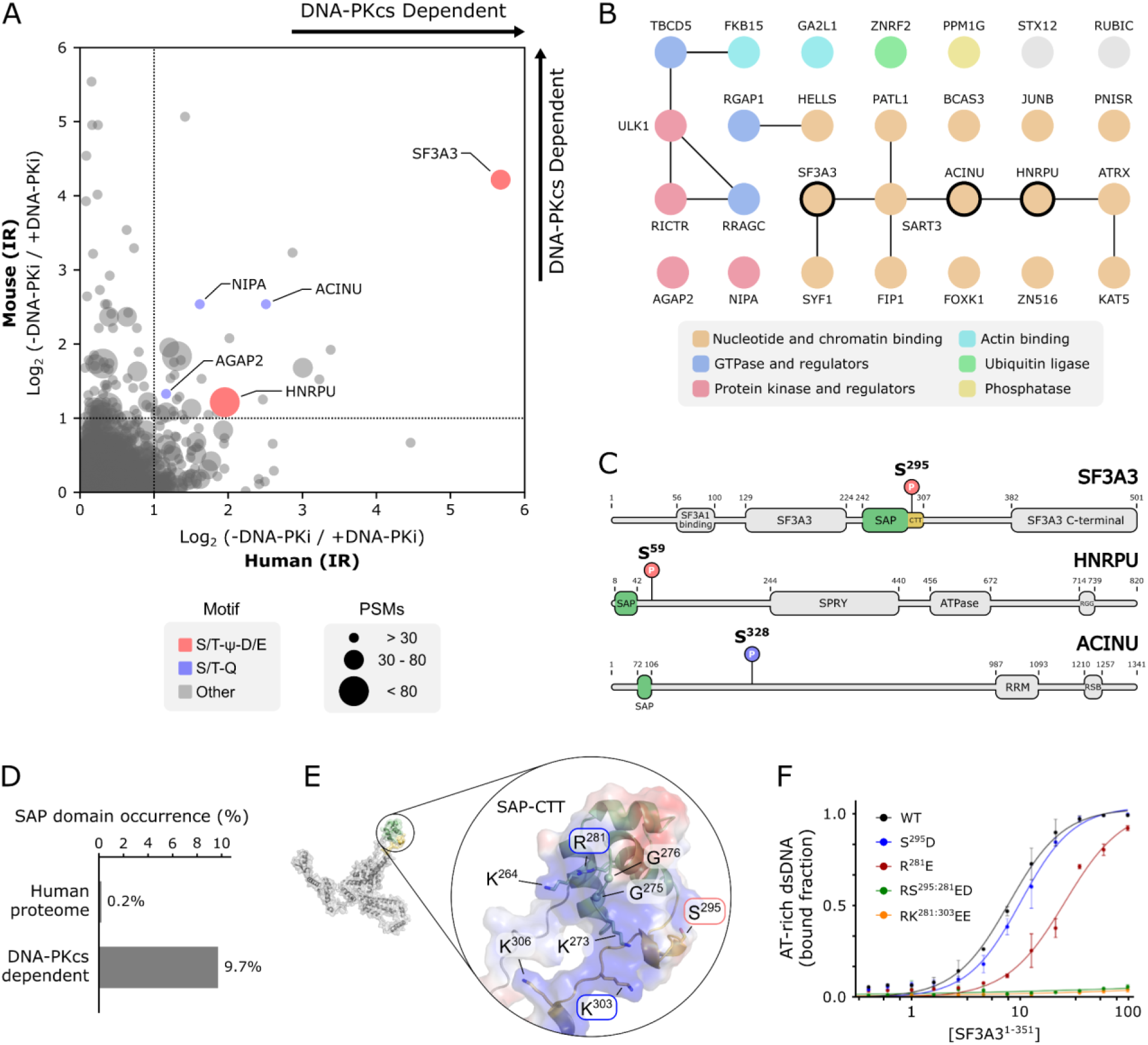
Cross-species analysis of DNA-PKcs-dependent phosphorylation defines a set of SAP-domain proteins as conserved targets. A) Scatter plot comparing IR-induced and DNA-PKcs-dependent phosphorylation sites found in both pre-B cells (mouse) and HCT116 cells (human). S/T-Q and S/T-ψ-D/E motifs are depicted in blue and red, respectively. The number of PSMs represents the combined count from both human and mouse phosphoproteomes. B) Cross-species proteins presenting conserved IR-induced and DNA-PKcs-dependent phosphorylation sites. Proteins containing the SAP domain are marked with a black circle. In the STRING network analysis, lines represent physical and/or functional interactions. C) Domain architecture of SAP-containing proteins, highlighting conserved IR-induced and DNA-PKcs-dependent phosphorylation at S/T-Q (blue) and S/T-ψ-D/E (red) motifs. D) The relative prevalence of the SAP domain in proteins with conserved IR-induced and DNA-PKcs-dependent phosphorylation sites (3 out of 31 proteins) compared to its occurrence in the Human proteome, as per reviewed sequences with an assigned SAP domain in the InterPro (https://www.ebi.ac.uk/interpro) database (23 out of 26,816 reviewed human proteins). E) Alphafold model of full-length human SF3A3. The inset zoom displays the electrostatic potential around the DNA binding interface of the SAP-CTT. Residues selected for charge reversal (R^281^E and K^303^E) and phosphomimetic (S^295^D) mutations are indicated in blue and red, respectively. The SAP domain and C-terminus tail (CTT) are illustrated in green and yellow, respectively. F) Plot showing fitted curves to the fraction of AT-rich dsDNA bound to increasing concentrations of wild-type (WT) and SF3A3^1-151^ variants (S^295^D, R^281^E, RS^295:281^ED, and RK^281:303^EE). Each data point represents the mean ± SD from three independent gel-shift assays (see Supp. Figure 5C).

### DNA-PKcs targets S/T-ψ-D/E motifs in SAP domain containing proteins

Most of the proteins found to be targeted by DNA-PKcs in both mouse and human cells were involved in splicing and transcription, and found to bind nucleotide molecules (Figure 5B). Kinases and phosphatases were also detected, consistent with the identification of DNA-PKcs-dependent phosphorylation at motifs other than S/T-Q or S/T-ψ-D/E, which could represent events controlled by kinases targeted by DNA-PKcs. Notably, several proteins containing DNA-PKcs-dependent phosphorylation at S/T-Q or S/T-ψ-D/E (potential direct targets) were RNA and/or DNA binding proteins (Figure 5B). Of these, three proteins (HNRPU, SF3A3 and ACINU) contained a SAP domain, which is an RNA/DNA-binding module typically present in proteins involved in DNA repair and RNA biology (Aravind & Koonin, 2000; Sahara *et al*, 1999; Inagawa *et al*, 2020; Tsuji *et al*, 2008). The presence of three SAP-domain proteins in the set of conserved DNA-PKcs-dependent phosphorylation events represents an approximately 50 fold enrichment over the frequency of these proteins in the whole proteome (Figure 5C). Notably, in the case of SF3A3 and HNRPU, the DNA-PKcs-dependent phosphorylation sites are located proximal to the SAP domain (Figure 5C), suggesting that phosphorylation of the S/T-ψ-D/E motifs may affect properties of the SAP domain. To investigate this hypothesis, we generated a soluble construct of the spliceosomal U2 snRNP protein SF3A3 (residues 1-351 - SF3A3^1-351^) and confirmed its ability to be phosphorylated by DNA-PKcs *in vitro* at serine 295, which is localized at a C-terminal tail extension (CTT) of the SAP domain (Supp. Figure 5A).

Next, we used electrophoretic mobility shift assay (EMSA) to investigate the DNA binding properties of SF3A3^1-351^ with an A-T-rich dsDNA oligonucleotide, known to be preferentially recognized by SAP domains (Renz & Fackelmayer, 1996). Structural predictions based on Alphafold SF3A3 models indicated that its SAP domain extends from the N-terminus SF3A1-interaction domain through flexible linkers (Figure 5E). Notably, the conserved S^295^ DNA-PKcs-dependent phosphosite within the CTT of the SAP domain is located within a hinge facing the canonical DNA-interacting interface (Figure 5E). Interestingly, the CTT is a structural feature uniquely conserved among SAP domains of SF3A3 and SDE2 homologs and has been implicated in DNA binding (Weinheimer *et al*, 2022). In line with the known affinity range observed for other SAP domains (Weinheimer *et al*, 2022), our EMSA showed that wild-type SF3A3^1-351^ binds dsDNA with an affinity of 7.9 ± 0.4 µM (Figure 5F and Supp. Figure 5D). Confirming that SF3A3 recognizes the A-T-rich dsDNA through its SAP domain, charge-reversal mutations at key residues on the predicted DNA-binding interface (R^281^E) and the CTT (K^303^E) decreased affinity by 3.3-fold (K_d_ = 26 ± 2 µM) for single R^281^E variant and entirely abolished DNA binding for the double RK^281:303^EE mutant SF3A3 (Figure 5F). Likewise, EMSA assays with a phospho-mimetic SF3A3 variant (S^295^D) revealed a reduced DNA-binding ability (K_d_ = 10.1 ± 0.5 µM), while the combination of S^295^D with the charge-reversal mutation at the SAP canonical DNA binding region (R^281^E) completely abrogated its dsDNA binding capacity (Figure 5F). This result suggests that the DNA-PKcs-mediated phosphorylation of SF3A3 at S^295^ S/T-ψ-D/E motif could influence its function by changing its ability to bind DNA.

## DISCUSSION

DNA-PKcs plays roles in multiple processes, including DNA repair, translation, and transcription (Shao *et al*, 2020; Caron *et al*, 2019; Jette & Lees-Miller, 2015). Despite well-established effects downstream of DNA-PKcs-mediated signaling being reported, the key substrates involved remain largely unknown. Mass spectrometry-based phosphoproteomics have enabled the mapping of signaling orchestrated by PIKK kinases, which has been mostly applied to the study of ATM and ATR signaling. Recent work has begun to map DNA-PKcs-signaling events (Schlam-Babayov *et al*, 2021), although the full scope of DNA-PKcs-dependent signaling remains incomplete given intrinsic issues with low coverage in phosphoproteomic analysis of complex protein mixtures. Here, to further expand the map of DNA-PKcs-dependent signaling, we used mouse pre-B cells lacking the Lig4 ligase, which accumulate unrepaired breaks and amplify DNA damage signaling, enabling increased coverage in the identification of DNA-PKcs signaling. This approach expanded the identification of DNA-PKcs-dependent signaling events and uncovered a novel motif preferentially targeted by DNA-PKcs, which was subsequently corroborated through *in vitro* kinase assays and cross-species analyses with human cells. The discovery of conserved features of DNA-PKcs kinase specificity opens new directions to investigate the mechanisms by which this kinase exerts its regulatory roles on DNA repair as well as on a multitude of other processes such as RNA processing and replication fork dynamics.

Our analysis identified over 2,000 phosphorylation sites induced by IR in G1 arrested pre-B cells and revealed that ATM and DNA-PKcs are responsible for the majority of the IR-induced signaling response in G1. Through the application of individual kinase inhibitors, we further revealed specific phosphorylation targets that rely exclusively on DNA-PKcs activity and are independent of ATM. Within this group of substrates, we observed an enrichment of the S/T-ψ-D/E motif in addition to the well characterized S/T-Q motif. Notably, the ability of DNA-PKcs to phosphorylate this motif in HNRPU (S^59^) and XRCC4 (S^260^) has been reported in low-throughput *in vitro* experiments (Berglund & Clarke, 2009; Yu *et al*, 2003), further corroborating our findings. Our study establishes the prevalence of the S/T-ψ-D/E motif *in vivo* and identifies several novel putative substrates phosphorylated at this motif. The finding of widespread substrate targeting at the S/T-ψ-D/E motif opens new directions to study DNA-PKcs signaling and the mechanisms of regulation of its substrates. It also raises the possibility that the preferential phosphorylation motif for DNA-PKcs may be regulated, switching from the conventional S/T-Q motif to the S/T-ψ-D/E motif depending on the context, such as differential localization or engagement with different protein complexes.

Analysis of DNA-PKcs-dependent phosphorylation sites conserved between mouse and human cells revealed the presence of several DNA/RNA binding proteins associated with splicing and transcription regulation. The group of proteins with conserved sites also included GTPase and kinase regulatory proteins, such as RICTR, RRAGC and AGAP2 notably involved in the crosstalk between DNA damage repair and pro-survival signaling pathways. For example, the Arf GAP protein AGAP2 is known to enhance PI3K activity, directly regulating the PI3K/Akt signaling cascade (Hu *et al*, 2005). Within the nucleus, AGAP2 interacts with PI3K in a GTP-dependent manner, stimulating its activity (Ye *et al*, 2000). The structural similarity of DNA-PKcs, a member of the PI3KK family, to PI3K and the S/T-Q phosphorylation site S927 observed in AGAP2 within our phosphoproteomes, suggests that AGAP2 may also interact with DNA-PKcs and modulate its activity. Our cross-species analysis also highlighted the enrichment of SAP domain-containing proteins amongst the group of IR-induced, DNA-PKcs-dependent sites. Although SAP domains are primarily recognized for their DNA-binding capability, studies have elucidated their involvement in RNA binding and protein-protein interactions as well (Payliss *et al*, 2022). The full extent of the relationship between SAP domain-containing proteins and DNA-PKcs remains unclear. Nonetheless, it is compelling to note that SAP domains are commonly observed in proteins involved in genome maintenance and DNA replication (Payliss *et al*, 2022; Weinheimer *et al*, 2022), two processes that are known to be regulated by DNA-PK. Moreover, KU70, a member of the DNA-PK holoenzyme that is crucial for its recruitment to DSBs and activation, also harbors a SAP domain (Inagawa *et al*, 2020). Given the high prevalence of SAP domain-containing proteins in the set of identified DNA-PKcs substrates, it is reasonable to hypothesize that these domains might facilitate the recruitment of substrate proteins to the proximity of active DNA-PKcs through protein-protein or protein-DNA interactions. It is tempting to speculate that SAP domains of KU70, SF3A3, HNRPU, and ACINU may all bind to a similar molecule/structure, which would explain how the substrate proteins get in proximity to DNA-PK. Moreover, the proximity of DNA-PKcs phosphorylation sites to the boundaries of SAP domains, such as those we identified in HNRPU (S^58^) and SF3A3 (S^295^), suggests that phosphorylation might modulate their affinity for binding partners. In fact, CDK1-induced phosphorylation of the SAP domain in SLX4 causes a conformational change from a flexible disordered loop to a more characteristic helix-loop-helix motif. This transition from disorder to order within this domain enhances its interaction with the structure-specific nuclease complex MUS81-EME1 and reduces its specificity for DNA substrates (Inagawa *et al*, 2020). Consistent with this finding, our *in vitro* binding assays using recombinant purified SF3A3^1-351^ and AT-rich dsDNA oligos demonstrated a reduced affinity of the phospho-mimetic variant, suggesting that phosphorylation at S^295^ might work cooperatively with the canonical SAP binding interface to modulate DNA interaction. Investigating the relationship between SAP domains and DNA-PKcs holds the potential to enhance our understanding of DNA-PKcs-specific signaling outcomes, many of which might not directly pertain to DNA repair.

Strikingly, the list of proteins we found to contain DNA-PKcs dependent phosphorylation at sites conserved between mouse and human did not include any of the proteins with established roles in DNA repair and DNA damage responses. We attribute this, in part, to the smaller coverage of the human dataset (since human cells had Lig4 and lower ratio of nuclear proteins) and the phosphorylation sites on DNA repair proteins tending to locate at residues or regions that are not well conserved. In fact, we did detect DNA repair and DNA damage response proteins with DNA-PKcs-dependent sites in both mouse and humans whose phosphorylation sites did not match conserved residues. This points to a limitation in our analysis and emphasizes the stringency of the list of 31 proteins presented in Figure 5B, which contains proteins with highly conserved sequences and that have a DNA-PKcs phosphorylation at a closely located residue in both organisms.

In conclusion, our findings uncover novel conserved features of DNA-PKcs kinase specificity and establish a connection between DNA-PKcs and SAP domain-containing proteins. Further exploration of these non-canonical modes of DNA-PKcs signaling holds the potential to address several outstanding questions, such as DNA-PKcs’s roles in regulating DNA repair-independent processes such as its role in the control of RNA associated processes and replication fork dynamics. The importance of comprehensively understanding DNA-PKcs signaling is further underscored by the use of DNA-PKcs inhibitors in clinical trials for cancer therapy (Matsumoto, 2022). For instance, our recent discovery that DNA-PKcs catalytic activity is essential for fork reversal expands the potential applications of DNA-PKcs inhibitors in combination therapies beyond their current use in conjunction with radiotherapy (Dibitetto *et al*, 2022). The use of DNA-PKcs inhibitors in rationalized combination therapies is particularly interesting given that these inhibitors are largely innocuous to non-cancer cells. Elucidation of novel DNA-PKcs substrates and signaling modalities will continue to be crucial for improving our understanding of DNA-PKcs biology and our ability to design improved therapeutic approaches.

## METHODS

### Mammalian Cell Culture and Cell Cycle Arrest

Mouse Pre-B cells (Lig4-/-) were a gift from Barry Sleckman and were grown at 37 ⁰C in Dulbecco’s modified Eagle medium (DMEM) supplemented with 10% Bovine Calf Serum (BCS), 1% non-essential amino acids, 0.0004% beta mercaptoethanol and penicillin/streptomycin solution (100 U/ml). Pre-B cell arrest in G1 was performed by treating 2 x 10^6^ cycling cells with 3μM imatinib for 48 hours. Cell cycle arrest was confirmed via flow cytometry by observing the loss of the populations representing S and G2/M phase cells after staining total DNA content with propidium iodide. Human HCT116 cells were purchased from ATCC and grown at 37 ⁰C in Dulbecco’s modified Eagle medium (DMEM) supplemented with 10% Bovine Calf Serum (BCS), 1% non-essential amino acids and penicillin/streptomycin solution (100 U/ml). Metabolic labeling was performed by growing cells for a minimum of five population doublings in SILAC medium supplemented with either “light” or “heavy” isotopes of lysine and arginine (Thermo).

### Induction of DNA Double Strand Breaks via Ionizing Radiation

Cells were exposed to 20 Gy of gamma radiation (0.89 Gy/min) from a Cs-137 source. The cell culture dishes were positioned at a designated location in the center of the chamber to ensure maximum dosage. They were continuously rotated to guarantee even distribution of the radiation across the entire sample. Cells were harvested 90 minutes after the treatment. In experiments involving kinase inhibition, cells were incubated at 37°C with 1 μM of ATM (KU-55933) and DNA-PKcs (NU7441) inhibitors for 1 hour prior to the ionizing radiation treatment.

### Protein Expression and Purification

For the chimeric polypeptides, 11-residues long sequences encoding ten S/T-ψ-D/E sites (Figure 4C), as identified in our phosphoproteomes, were codon-optimized for *Escherichia coli* protein expression, and arranged back-to-back to create synthetic chimeric peptide oligonucleotides, both wild-type and positional mutants (S/T-A-D/E and S/T-ψ-A). These synthetic oligonucleotides were synthesized and cloned into a bacterial expression vector based on pET28a (Novagen), which adds a cleavable 6xHIS-SUMO tag at the N-terminal (pETSUMO). A construct encompassing residues 1-351 of human SF3A3 (SF3A3^1-351^) was amplified from a plasmid containing the full-length ORF sequence of human SF3A3 using the primers 5’-GAGAACAGATTGGTGGAATGGAGACAATA and 5’-GGAGCTCGAATTCGGATTACCTGGCTTGCTTGCGCTG-3’, and cloned into pETSUMO. The chimeric polypeptides and SF3A3^1-351^ were overexpressed in *E. coli* BL21 (DE3). The cultures were grown at a temperature of 37 °C in LB medium, supplemented with 50 μg/ml kanamycin. Upon reaching an optical density at 600 nm (OD600) of ∼0.6, protein expression was induced by the addition of 0.3 mM IPTG, and allowed to proceed for 16 hours at a temperature of 18 °C. Following this, the cells were collected via centrifugation and resuspended in lysis buffer (25 mM Hepes, 300 mM NaCl, 20 mM imidazole, pH 8.0). After lysis through sonication, clear lysates were extracted through centrifugation (20,000 xg), and then loaded onto columns containing 1 mL of pre-equilibrated Co-NTA resin (Malvergen) with lysis buffer. The resin was washed with ten column volumes (CV) of lysis buffer and followed by an additional washing step with ten CV of buffer A (25 mM Hepes, 300 mM NaCl, pH 8.0). Protein elution was then conducted using five CV of buffer B (25 mM Hepes, 300 mM NaCl, 300 mM imidazole, pH 8.0). The 6xHis-SUMO moiety of eluted proteins was removed by specific cleavage using the yeast protease Ulp-1. Finally, the cleaved proteins underwent size exclusion chromatography using a Superdex 75 column (Cytiva) that was pre-equilibrated with a buffer consisting of 25 mM Hepes at pH 8.0 and 150 mM NaCl. This step was crucial for separating the expressed proteins from the cleaved fusion tags and Ulp-1. Purified proteins were concentrated on Cytiva filters (10 KDa cutoff) to ∼10 mg/ml and stored at −80 °C.

### In vitro Kinase Assay

The assays were conducted using the DNA-PK Kinase Enzyme System (Cat # V4106, Promega), following the provided instructions. Briefly, reaction mixtures were composed of 10 μg of chimeric polypeptide, 0.1 mg/mL BSA, 1X activation buffer (10 μg/mL calf thymus DNA), 1 mM DTT, 1 X reaction buffer, 50 units of DNA-PK, and 150 μM ATP, in a total volume of 50 μL. The reactions were initiated by adding ATP, followed by incubation at room temperature for 1 h. Once completed, the reactions were subjected to a 5-minute incubation at 95 °C to inactivate the DNA-PK, flash-frozen in liquid nitrogen and preserved at -80 °C.

### Electrophoretic Mobility Shift Assay (EMSA)

DNA oligonucleotides (5′-[5IRD700]-GAAAAAATATAAAATAGCTAGTTTTATTTTATTATTT TCTGAAT-3’ and 5′-[5IRD700]-AATTCAGAAAATAATAAAATAAAACTAGCTATTTTATAT TTTTTC-3’) were dissolved in annealing buffer (10 mM Tris, 100 mM KCl, pH 7.5) to a final concentration of 20 µM. The oligo strands were mixed in equimolar amounts and annealing was performed by heating to 95 °C and slowly cooling to room temperature. Binding reactions were performed with increasing protein amounts (0 – 200 µM) and 2 nM oligonucleotides duplex in binding buffer (50 mM Tris, 5% glycerol, 150 mM KCl, and 1 mM DTT) for 30 min at room temperature. Samples were loaded onto polyacrylamide native gels (5% polyacrylamide 29:1 [BioRad], 1× TBE, 0.3% APS, and 0.06% Temed) and protein/DNA complexes were resolved by electrophoresis at 60 V for 2 h at 24 °C in 1X TBE. Gels were imaged using a Li-Cor Odyssey scanner and bands were quantified using Image Studio 4.0 software (LI-COR Biosciences).

### Sample Preparation for Phosphoproteomic Analysis

After treatment Pre-B cells were harvested by centrifugation at 1000 xg for 5 minutes. HCT116 cells were detached from cell culture plates with 0.25% Trypsin (Thermo, Cat: 25200056). The trypsin was deactivated by adding 4X volume of fully supplemented DMEM then cells were pelleted by centrifugation at 1000 xg. Harvested cell pellets were washed once with cold PBS followed then lysed with cold modified RIPA buffer (50mM Tris-HCl, pH 7.5, 150mM NaCl, 1% Tergitol, 5mM EDTA supplemented with complete EDTA-free protease inhibitor cocktail (Roche) and PhosSTOP (Millipore)) for 30 minutes on ice. 4mg of light and heavy lysates were mixed then reduced and denatured with 1% SDS and 5mM DTT, respectively at 42 ⁰C. Cysteines of the denatured proteins were alkylated by incubation with 25mM iodoacetamide for 15 minutes in the dark. Lysates were then precipitated with a cold precipitation solution (50% acetone. 49.9% ethanol, 0.1% acetic acid). After precipitation, the protein pellet was solubilized with 2M urea then digested with TPCK-trypsin overnight at 37⁰C. Digested peptides were then desalted using a Waters 20mg Sep-pak C18 column then dried in a SpeedVac and resuspended in 1% acetic acid. Enrichment of phosphopeptides was accomplished using High Select Fe NTA phosphopeptide enrichment kit. Enriched phosphopeptides were then dried and resuspended in 15uL H2O and 10uL 10% formic acid.

### HILIC Fractionation

Immediately before injection 60 μL of HPLC grade ACN was added to the phosphopeptide samples. Samples were then fractionated via hydrophilic interaction liquid chromatography. A gradient was generated using three buffers: Buffer A (90% ACN), Buffer B (75% ACN and 0.0005% TFA), and Buffer C (0.025% TFA). The gradient consisted of 100% Buffer A at 0 minutes, 88% Buffer B and 12% Buffer C at 5 minutes, 60% Buffer B and 40% Buffer C at 30 minutes, and 5% Buffer B and 95% Buffer C from 35 to 40 minutes. Fractions were collected every 60 seconds between minutes 8 and 34. Fractions were then dried in a speedvac and resuspended in 0.1% TFA.

### Proteomic Data Acquisition

HILIC fractions were analyzed using liquid chromatography-tandem mass spectrometry (LC-MS/MS) with a Q Exactive HF instrument. Peptides were separated using an UltiMate 3000 RSLC nano chromatographic system. The column used was 30 cm long with an inner diameter of 100 μm, packed with Reprosil Pur C18AQ 3 μm resin. Data acquisition was performed with Xcalibur software from Thermo Fisher Scientific, and the Q-Exactive was operated in data-dependent mode. Survey scans were conducted in the Orbitrap mass analyzer, covering the mass range of 380 to 2000 m/z, with a mass resolution of 60,000 (at m/z 200). MS/MS analysis was performed by selecting the most abundant ions with a charge state of 2, 3, or 4 within an isolation window of 2.0 m/z. The selected ions were fragmented using Higher-energy Collisional Dissociation (HCD) with a normalized collision energy of 28, and the tandem mass spectra were acquired in the Orbitrap mass analyzer with a mass resolution of 15,000 (at m/z 200).

For the Pre-B and HCT116 phosphoproteomic experiments, the Uniprot databases for mouse and human, respectively, were utilized. Peptide identification and quantification were processed using the trans proteomic pipeline (TPP) tools (Deutsch et al. 2010). The search engine employed was Comet (v. 2019.01.1) (Eng, Jahan, and Hoopmann 2013). Search parameters included a requirement for semi-tryptic peptides, a precursor match tolerance of 15 ppm, differential mass modification of 79.966331 Da for phosphorylated peptides, and a static modification of 57.021465 Da for alkylated cysteine residues. After the searches, peptides were scored using the PeptideProphet algorithm, and SILAC ratios were calculated using XPRESS. Resultant datasets were filtered using the following parameters: minimum probability of 0.9, minimum peptide length of 7 amino acid residues, accurate mass binning, and restriction to +2, +3, and +4 ion charge states (Keller et al. 2002). Phosphorylated peptides were further evaluated using PTMProphet to obtain a localization score for the modification (Shteynberg et al. 2019).

### Phosphorylation Motif Analysis

Phosphorylation events were categorized into several motif categories based on the residues in the +1 and +2. Residues in each position were separated into several categories. Ψ indicates the bulky hydrophobic residues phenylalanine, isoleucine, leucine, and valine. D/E represents the negatively charged residues aspartic acid and glutamic acid glutamic acid. K/R represents the positive residues lysine and arginine. P and Q represent proline and glutamine, respectively. S/T/Y represents the phosphorylatable residues serine, threonine and tyrosine. For each experiment (e.g. IR + DNA-PK inhibitor) the proportion of each motif among the induced and kinase dependent sites were calculated using the ratio of the number of sites to all of the sites in that. These proportions were used to generate matrices that could be converted to heatmaps using python.

### Data Availability

Mass spectrometry data generated from this study has been deposited to the PRIDE database under the identifier: PXD042258 (https://www.ebi.ac.uk/pride/).

## ACKNOWLEDGEMENTS

We thank Beatriz S. Almeida for technical support and members of the Smolka Lab for valuable discussions. This work is supported by a grant from the National Institute of Health, R35GM141159 to M.B.S.

---

**Supp. Figure 1.**
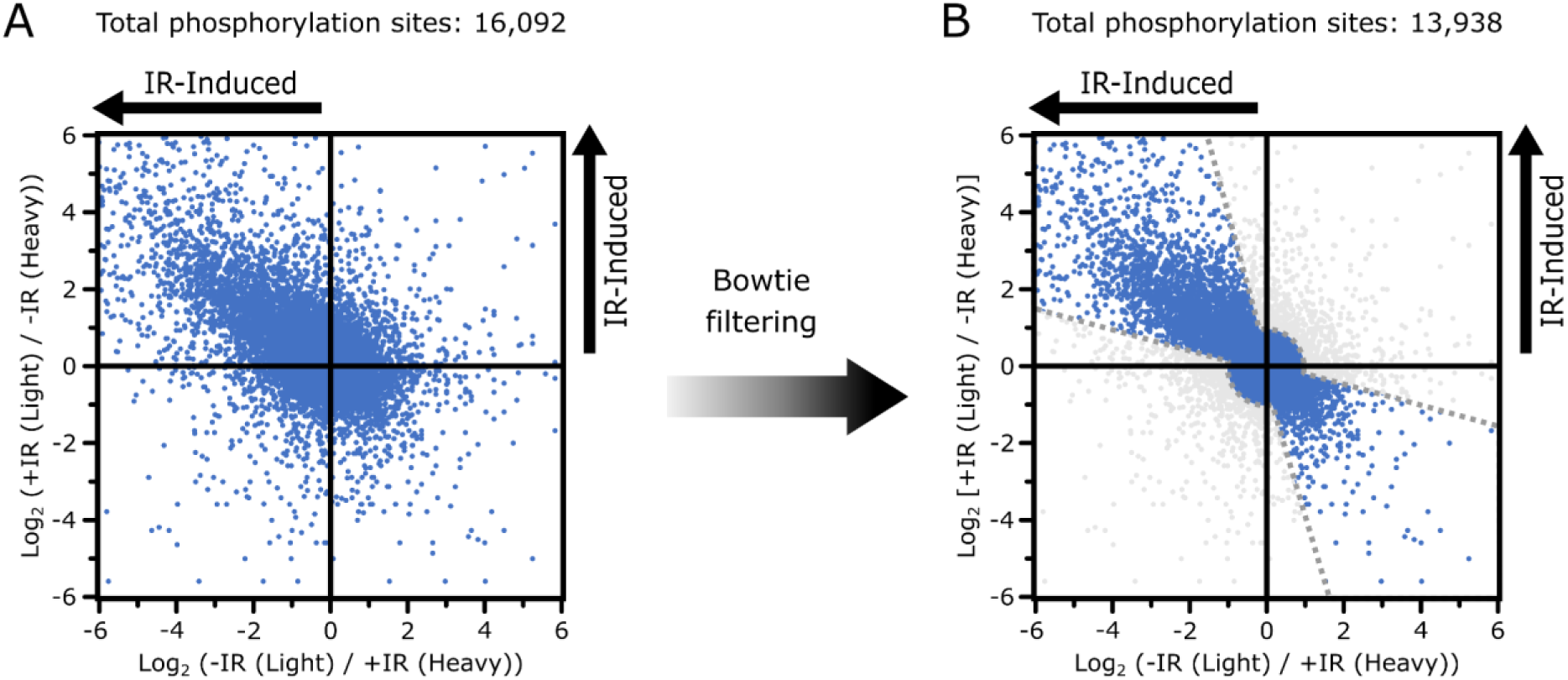
Phosphoproteomics of IR-treated Mouse pre-B cells. A) Scatter plot comparing all phosphosites identified in two quantitative phosphoproteomic analyses of cells treated (+IR) and untreated (-IR) with ionizing radiation. The isotope labeling regimen (Light/Heavy) is reversed in the two experiments. B) Following Bowtie filtering (Faca et al., 2020), the number of high-confidence sites, indicated by light blue dots within the dashed gray lines, decreased from 16,092 to 13,938.

**Supp. Figure 2.**
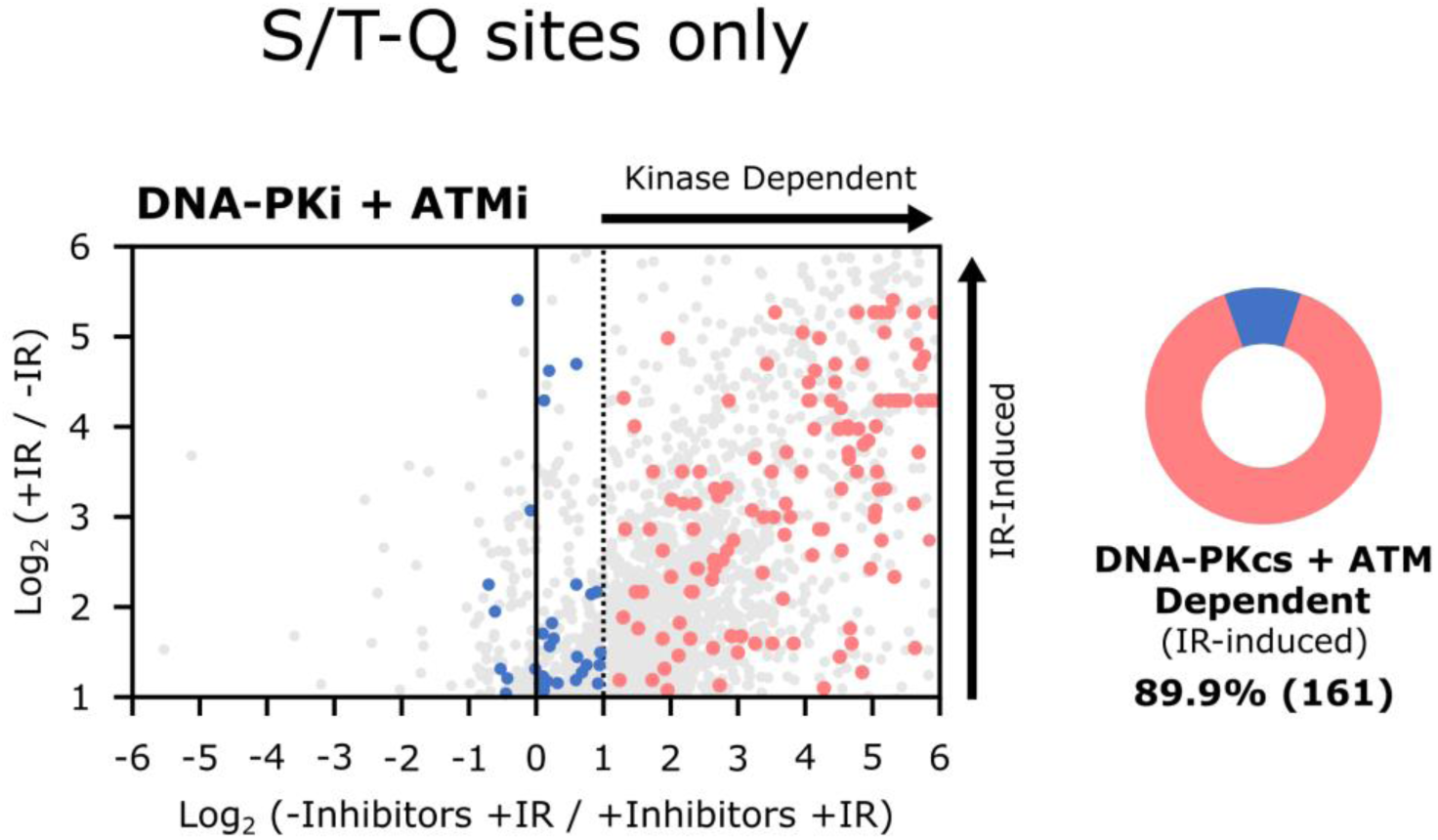
Enrichment of IR-induced S/T-Q sites dependent on DNA-PKcs and ATM activities. This figure reproduces the scatter plot from Figure 2C, highlighting IR-induced S/T-Q sites that are dependent (in light red) and independent (in blue) on the activities of DNA-PKcs and ATM.

**Supp. Figure 3.**
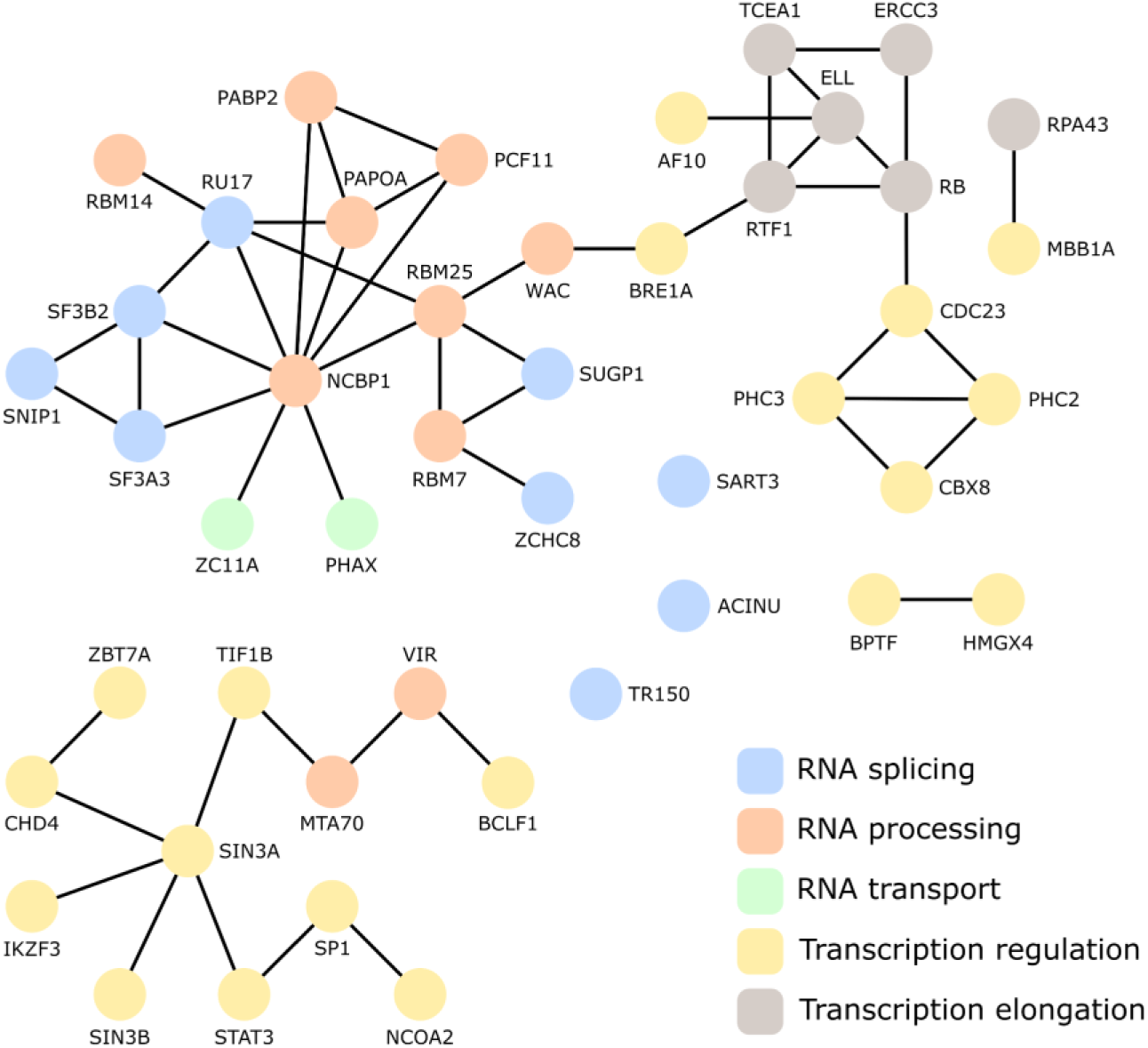
DNA-PKcs and ATM regulate the phosphorylation status of various proteins involved in RNA biology. String analysis of proteins exhibiting phosphorylation sites downregulated by independent treatment with ATM and DNAPKcs inhibitors.

**Supp. Figure 4.**
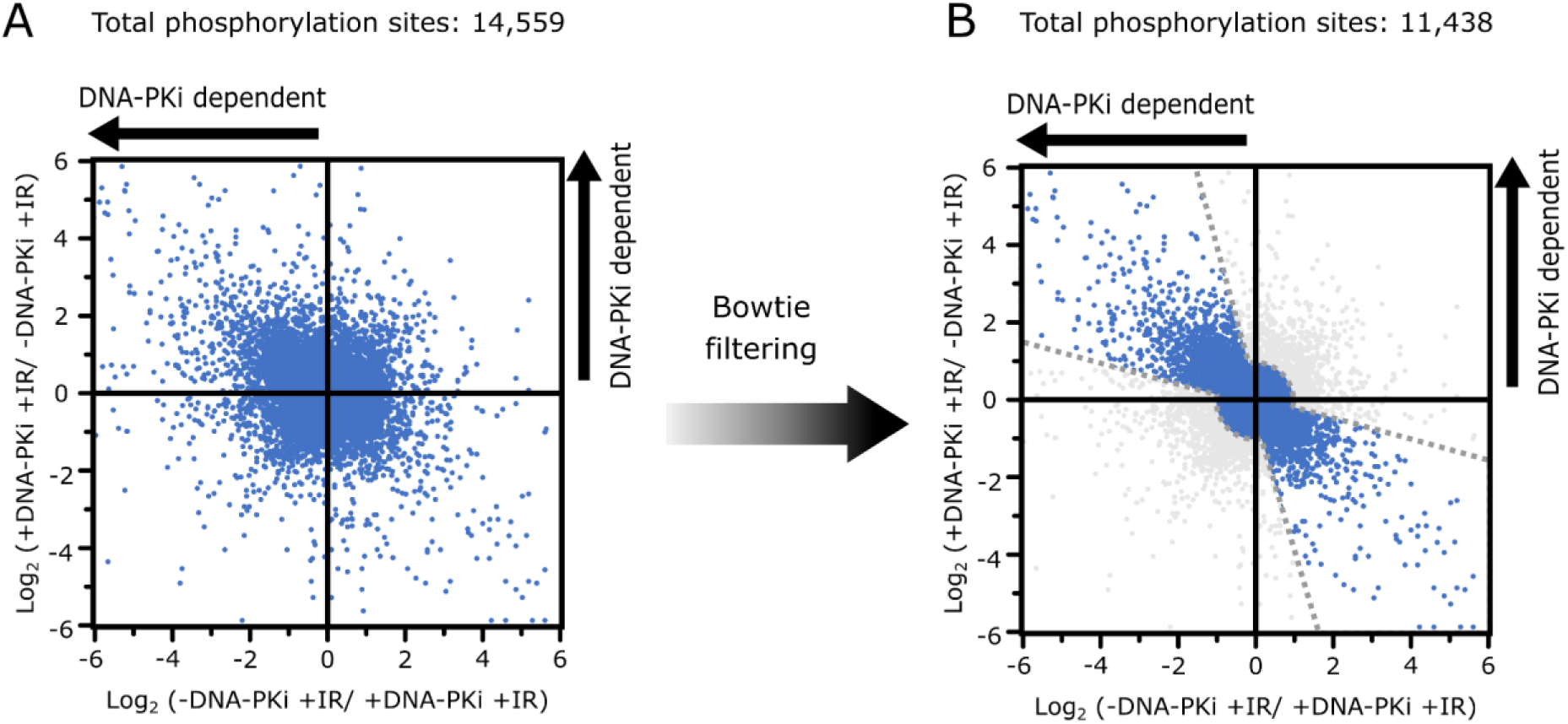
Phosphoproteomics of IR-treated Human HCT116 cells in the presence or absence of DNA-PKcs inhibitor. A) Scatter plot comparing all phosphosites identified in two quantitative phosphoproteomes of IR irradiated cells, one set pre-treated with DNA-PKcs inhibitor (+DNA-PKi) and the other untreated (-DNA-PKi). The isotope labeling regimen (Light/Heavy) is reversed in the two experiments. B) After Bowtie filtering (Faca et al. 2020), the number of high confidence sites, indicated by light blue dots within the dashed gray lines, reduced from 14,559 to 11,438.

**Supp. Figure 5.**
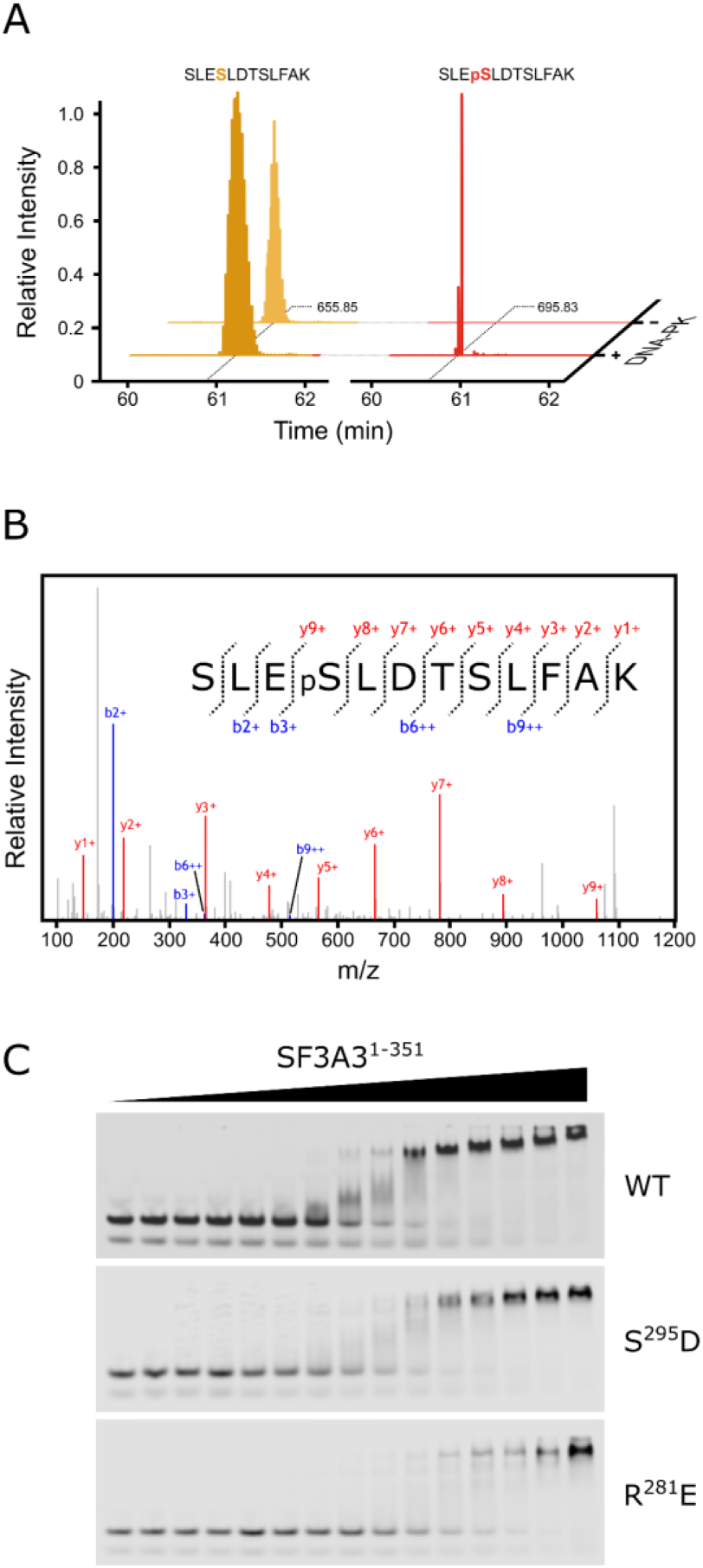
*In vitro* kinase assay and EMSA analysis performed with recombinant SF3A3^1-351^. A) Reconstructed ion chromatogram of the SF3A3 peptide containing S^295^ (SLES^295^LDTSLFAK) demonstrates that the phosphorylated form is only detected following *in vitro* phosphorylation by DNA-PK. B) MS2 spectrum confirms the identification of the SLEpS^295^LDTSLFAK phosphopeptide. C) EMSA results of AT-rich dsDNA labeled with near-infrared fluorescent dye incubated with increasing concentrations of SF3A3^1-351^ WT and variants (S^295^D and R^281^E).

